# Deep brain stimulation of the nucleus accumbens shell does not decrease cocaine self-administration in cocaine-dependent rats but increases GluR1/GluA1 in the central nucleus of the amygdala

**DOI:** 10.1101/438507

**Authors:** Jenni Kononoff, Philippe A. Melas, Johanna S. Qvist, Giordano de Guglielmo, Marsida Kallupi, Eric R. Kandel, Olivier George

**Author notes:** Corresponding author: Dr. Olivier George, Department of Neuroscience, The Scripps Research Institute, 10550 North Torrey Pines Road, SP30-2400, La Jolla, CA 92037, USA.

## Abstract

**Background:** Cocaine addiction is a major public health problem. Despite decades of intense research, no effective treatments are available. Both preclinical and clinical studies of drug addiction strongly suggest that the nucleus accumbens (NAcc) is a viable target for deep brain stimulation (DBS).

**Objective:** Although previous studies have shown that DBS of the NAcc decreases cocaine seeking and reinstatement, the effects of DBS on cocaine intake in cocaine-dependent animals have not yet been investigated.

**Methods:** Rats were made cocaine-dependent by allowing them to self-administer cocaine in long-access sessions (6 h, 0.5 mg/kg/infusion). The effects of high-frequency DBS of the NAcc shell on cocaine intake was then studied. Furthermore, cocaine-induced locomotor activity, irritability-like behavior during cocaine abstinence and the levels of the α-amino-3-hydroxy-5-methyl-4-isoxazolepropionic acid (AMPA) receptor subunits 1 and 2 (GluR1/GluA1 and GluR2/GluA2) after DBS were investigated.

**Results:** Contrary to our expectations, DBS of the NAcc shell induced a slight increase in both cocaine self-administration and cocaine-induced locomotor activity. In addition, 18 h into cocaine withdrawal, we found that DBS decreased irritability-like behavior. We also found that DBS-induced a robust increase in both cytosolic and synaptosomal levels of GluR1, but not GluR2, specifically in the central nucleus of the amygdala but not in other brain regions.

**Conclusions:** These preclinical results with cocaine-dependent animals do not support high-frequency DBS of the NAcc shell as a therapeutic approach for the treatment of cocaine addiction in active cocaine users. However, the decrease in irritability-like behavior during cocaine abstinence, together with previous findings showing that DBS of the NAcc shell reduces the reinstatement of cocaine seeking in abstinent animals, warrants future investigations of DBS as a treatment for negative emotional states and craving during abstinence.

**Highlights:**

- High-frequency DBS of the NAcc shell for the treatment of cocaine addiction is proposed
- DBS of the NAcc shell does not decrease cocaine intake in cocaine-dependent rats
- DBS increases the level of GluR1 specifically in the central nucleus of the amygdala

## Introduction

Cocaine use disorder is a global public health issue. In 2016 alone, the number of new cocaine users in the United States increased to 1.1 million [1]. However, despite decades of intense research, no effective treatments for cocaine addiction are available [2], highlighting the critical need for effective alternative therapeutic strategies.

Based on its 30-year history of success for the treatment of movement disorders, deep brain stimulation (DBS) may be a promising strategy for the treatment of severe substance use disorder. To date, nine clinical studies have been published (with a total of only 25 participants) that studied the effects of DBS of the nucleus accumbens (NAcc) for the treatment of substance use disorder. These studies reported a reduction of craving or consumption of alcohol [3–5], tobacco [6, 7], opioids [8–10], and cocaine [11]. However, given the low number of subjects in these studies (including a single case study for cocaine) and the well known publication bias toward positive results [12–14], the results of these clinical studies need to be interpreted with caution [15].

In addition to clinical investigations, several animal studies have demonstrated that DBS of the NAcc decreases addiction-related behaviors [16]. For example, DBS of the NAcc has been found to prevent morphine-induced conditioned place preference [17], reduce methamphetamine seeking and intake [18], decrease alcohol drinking [19, 20], and decrease the reinstatement of cocaine seeking [21, 22]. Although these studies are very promising, they did not directly test the effects of DBS of the NAcc on excessive cocaine intake in dependent animals. This is a critical step before considering translating the DBS approach to humans.

In the present study, we investigated the effects of DBS of the NAcc shell on the escalation of cocaine self-administration, locomotor sensitization, and irritability-like behavior in cocaine-dependent rats that were given extended access (6 h/day) to cocaine. We further evaluated the effects of DBS of the NAcc on key brain regions (NAcc shell, NAcc core, dorsal striatum, ventral prefrontal cortex [vPFC], dorsal PFC [dPFC], central nucleus of the amygdala [CeA], insular cortex [insula], and ventral tegmental area [VTA]) using the levels of glutamate receptors GluR1 and GluR2 as a proxy for glutamatergic transmission. The dynamic regulation of GluR1/GluR2 during cocaine self-administration has been suggested to be a critical factor in the transition to cocaine addiction [23–27].

## Materials and Methods

### Animals

For the experimental design, see Fig. 1A. Briefly, male Sprague Dawley rats (Charles River, Wilmington, MA, USA; *n* = 14) were first trained to self-administer cocaine for 1 h per day for 14 sessions. They then underwent daily 6-h sessions for 17 sessions. The effects of DBS were tested after stabilization of the escalation of cocaine intake. The rats were housed two per cage under a reverse 12 h/12 h light/dark cycle in a temperature (20-22°C)- and humidity (45-55%)-controlled animal facility with *ad libitum* access to food and water. All of the procedures were conducted in adherence to the National Institutes of Health Guide for the Care and Use of Laboratory Animals and were approved by the Institutional Animal Care and Use Committee of The Scripps Research Institute.

**Figure 1.**
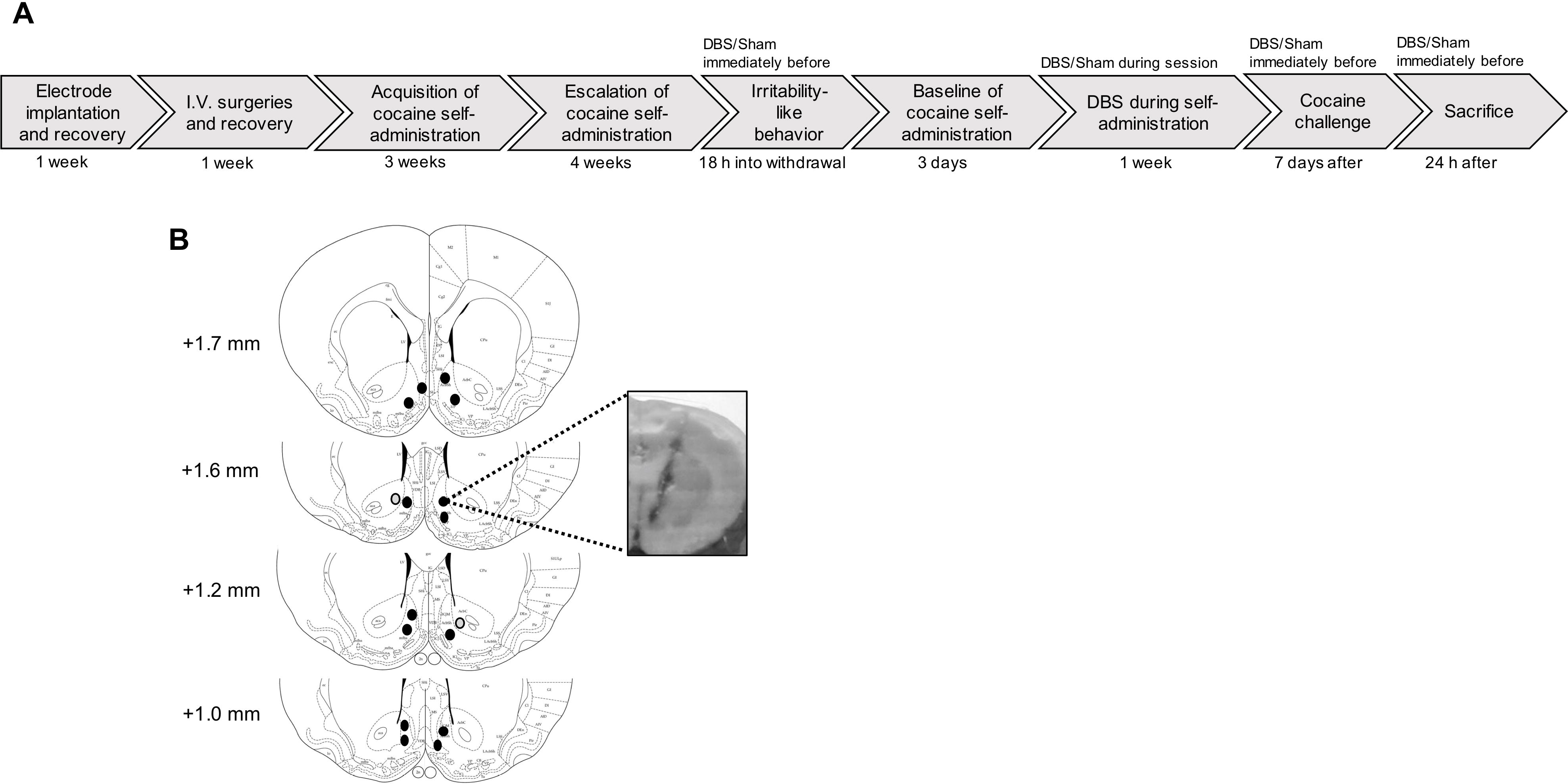
Experimental design and electrode placements. (A) Timeline of the experiment. (B) Schematic illustration of electrode placements in the NAcc shell (black dots) and a representative image of the placement. The gray dots represent misplaced electrodes.

### Drugs

Cocaine HCl (National Institute on Drug Abuse, Bethesda, MD, USA) was dissolved in 0.9% saline (Hospira, Lake Forest, IL, USA) at a dose of 0.5 mg/kg/0.1 ml infusion and self-administered intravenously. For the cocaine challenge experiment, cocaine was dissolved in 0.9% saline at a dose of 10 mg/kg and was injected intraperitoneally (i.p).

### Electrode implantation

The animals were anesthetized and mounted on a stereotaxic apparatus (Kopf Instruments, Tujunga, CA, USA). Bipolar stainless-steel electrodes (MS303/3, Plastics One, Wallingford, CT, USA) were implanted in the NAcc shell using the following coordinates with reference to bregma: 17° angle toward midline; anterior/posterior, +1.0 mm; medial/lateral, ±3.0 mm; dorsal/ventral, −7.0 mm from dura. The coordinates were selected based on a previous study [22] using the Paxinos and Watson rat brain atlas [28]. The electrodes were fastened to the skull with dental acrylic with four stainless-steel screws. The electrode placement was verified from fresh-frozen sections. One rat was excluded because of misplaced electrodes, leaving *n* = 6 rats in the Sham group and *n* = 7 rats in the DBS group for the final behavioral analysis.

### Intravenous catheterization

Once the animals recovered from electrode implantation, they were allowed 1 week to recover from surgery. They were anesthetized, and intravenous catheters were aseptically inserted in the right jugular vein using a modified version of a procedure that was described previously [29, 30]. The catheters were flushed daily with heparinized saline (10 U/ml of heparin sodium; American Pharmaceutical Partners, Schaumburg, IL, USA) in 0.9% bacteriostatic sodium chloride that contained 20 mg/0.2 ml of the antibiotic Cefazolin (Hospira, Lake Forest, IL, USA).

### Operant training

Self-administration was performed in operant chambers (Med Associates, St. Albans, VT, USA). Each chamber was equipped with two retractable levers. Cocaine was delivered through an infusion pump that was activated by responses on the right (active) lever, resulting in the delivery of cocaine (0.5 mg/kg/0.1 ml). Responses on the left (inactive) lever were recorded but had no scheduled consequences. The rats were first trained to self-administer cocaine under a fixed-ratio 1 (FR1) schedule of reinforcement in daily 1-h sessions until a stable baseline of reinforcement was achieved (± 10% variation over the last three sessions). A 20-s timeout (TO) period followed each cocaine infusion. After the acquisition period, the rats were subjected to daily 6-h cocaine self-administration sessions to allow them to escalate their cocaine intake [31].

### Deep brain stimulation

The DBS and sham treatments were conducted in the operant self-administration chambers. Before starting the experiments, the rats were habituated to being attached to the stimulating cables at least five times. A swivel (Plastics One, Roanoke, VA, USA) facilitated the cable connections between the head cap and stimulation system to allow free movement. A digital stimulator (model DS8000, World Precision Instruments, (Sarasota, FL, USA) was connected to a digital stimulus isolator (model DLS100, World Precision Instruments, Sarasota, FL, USA). The DS8000 stimulator was set to 1.5 V, and the DLS100 stimulus isolator was set to 1 mA to produce 150 μA stimulation. Each stimulator was checked with an oscilloscope to ensure correct stimulation amplitude and frequency. For DBS, we used bilateral, monophasic (60 μs pulse width and 130 Hz frequency) stimulation with a constant current of 150 μA. These parameters are consistent with previous studies in the field that used high-frequency stimulation (HFS; [21, 32–34]. After a stable level of self-administration was achieved, the effects of DBS on cocaine self-administration in cocaine-dependent rats was evaluated for 4 consecutive days, with 2 “ON” and 2 “OFF” sessions on alternate days. During the “ON” session in the DBS group, the stimulation was turned on 30 min before the levers were extended for self-administration and lasted 90 min during the self-administration session (total of 2 h). For the remaining 4.5 h of the 6-h self-administration session, the animals were connected to the stimulating cables but not stimulated. The rats in the Sham group were connected to the stimulating cables during the “ON” session but were not stimulated. During the “OFF” sessions, the rats were not connected to the stimulating cables in either the DBS or Sham group. For the subsequent behavioral experiments (irritability-like behavior and locomotor activity) and before euthanizing the rats to collect brains, the DBS was “ON” for 60 min.

### Irritability-like behavior

All behavioral testing was conducted during the dark phase. To test irritability-like behavior, we used the bottle-brush test, based on the experimental method that was designed for mice [35, 36] and slightly modified by our laboratory [37]. Briefly, the animals were randomized, and three observers scored the behaviors. Testing consisted of 10 trials with 10-s intertrial intervals in plastic cages (27 cm × 48 cm × 20 cm). A bottle-brush was rotated rapidly toward the rat’s whiskers. Both aggressive responses (smelling, biting, boxing, following, and exploring the brush) and defensive responses (escaping, digging, jumping, climbing, defecation, vocalization, and grooming) were recorded. The aggressive and defensive responses were chosen based on the methods of Riittinen et al. (1986) and Lagerspetz and Portin (1968). Both aggressive and defensive behaviors were summed to calculate the total irritability score. Irritability-like behavior was assessed 18 h into withdrawal once the rats were made dependent on cocaine, and the rats were subjected to DBS or sham treatment for 60 min immediately before testing.

### Locomotor activity after cocaine challenge

Seven days after the last cocaine self-administration session, the rats were challenged with cocaine (10 mg/kg, i.p.), and locomotor activity was measured. The animals were first allowed to habituate to the test chamber (43.2 cm × 43.2 cm × 30.5 cm, Med Associates, St. Albans, VT, USA) for 30 min. The rats were then injected with cocaine, and locomotor activity was monitored for 60 min under red light. Locomotor activity (in centimeters) was recorded using a video camera that was connected to the ANY-maze Video Tracking System 5.11 (Wood Dale, IL, USA).

### Tissue preparation

Twenty-four hours after the cocaine challenge, rats were subjected to either DBS or sham treatment for 60 min immediately before they were euthanized. The brains were removed, frozen on dry ice, and kept at −80°C until punched. The brains were blocked in the coronal plane and sectioned at 500 μm thickness using a cryostat. The NAcc shell and core, dorsal striatum, vPFC, dPFC, CeA, insula, and VTA were punched based on a previously described technique [38] with modifications. The punch sites for each region were determined based on the rat brain atlas [28].

### Western blot

α-Amino-3-hydroxy-5-methyl-4-isoxazolepropionic acid (AMPA) receptors exist both at synapses and in internal stores and undergo trafficking [39–41]. Thus, in the present study, we measured both synaptosomal and cytosolic levels of GluR1 and GluR2. Brain samples were processed using Syn-PER Synaptic Protein Extraction Reagent (ThermoFischer Scientific, Waltham, MA, USA), which allows the separation of synaptosomal and cytosolic neuronal fractions from the same samples. Protein concentration measurements were performed using the Pierce Detergent Compatible Bradford Assay Kit (Thermo Scientific), and equal amounts of sample were run on 4-20% Mini-PROTEAN TGX Precast Protein Gels (Bio-Rad Laboratories, Hercules, CA, USA) and then transferred to Immobilon-FL polyvinylidene difluoride membranes (EMD Millipore, Billerica, MA, USA). Following incubation with primary and secondary antibodies and appropriate washes, protein bands of interest were visualized using a fluorescent detection system (Odyssey Classic, LI-COR Biotechnology, Lincoln, NE, USA). The membranes were stripped and re-probed using Restore Fluorescent Western Blot Stripping Buffer (Thermo Scientific, Waltham, MA, USA) between target and normalization antibody incubations. Primary antibodies were used against the AMPA receptor subunits GluR1/GluA1 (catalog no. ab109450, Abcam, Cambridge, MA, USA) and GluR2/GluA2 (catalog no. 13607, Cell Signaling, Danvers, MA, USA, or catalog no. sc517265, Santa Cruz Biotechnology, Santa Cruz, CA, USA) and the normalization controls synaptophysin (catalog no. 5461, Cell Signaling, Danvers, MA, USA) and β-actin (catalog no. ab6276, Abcam, Cambridge, MA, USA). Protein bands were quantified using Image Studio software (LI-COR). Synaptosomal fractions were normalized to synaptophysin, and cytosolic fractions were normalized to β-actin. The data were transformed to fold-change differences relative to the control (Sham) group.

### Statistical analyses

The escalation of cocaine self-administration was analyzed using one-way repeated-measures analysis of variance (ANOVA) or Student’s paired *t*-test (two-tailed). The effect of DBS and sham treatments on the escalation of cocaine self-administration was analyzed using either two-way repeated-measures ANOVA or Student’s paired *t*-test (two-tailed) using the average intake in the two OFF/ON sessions. The effect of DBS on irritability-like behavior was analyzed using unpaired *t*-test (two-tailed). Cocaine-induced locomotor activity was analyzed using two-way repeated-measures ANOVA. Differences in protein levels of GluR1 and GluR2 were analyzed using two-way ANOVA followed by Bonferroni’s multiple-comparison *post hoc* test. The Western blot data were analyzed using two-way ANOVA followed by Bonferroni’s multiple-comparison *post hoc* test. Values of *p* < 0.05 were considered statistically significant. All of the analyses were performed using Prism 7 software (GraphPad, La Jolla, CA, USA).

## Results

### Escalation of cocaine self-administration

Our experimental design is illustrated in Fig. 1A. The electrode placements are shown in Fig. 1B. The rats were trained to self-administer cocaine during daily 1-h sessions. After the acquisition period (Fig. 2A), the rats were allowed to escalate their cocaine intake during daily 6-h sessions (Fig. 2B). The one-way repeated-measures ANOVA (*F*_16,192_ = 6.597, *p* < 0.0001) showed that the rats significantly increased their intake starting in session 9 (Fig. 2B; Bonferroni’s *post hoc* comparison with baseline: *p* < 0.05 for sessions 9 and 10, *p* < 0.01 for session 11, and *p* < 0.0001 for sessions 12-17). Fig. 2C shows the significant escalation of cocaine intake during the first hour of the 6-h session (i.e., the loading phase) between session 1 and session 17 (*t*_12_ = 4.54, *p* = 0.0007). For the DBS studies, the animals were divided into two groups, with no significant difference between the Sham and DBS groups in the average cocaine intake during the last 5 sessions (*t*_11_ = 0.2983, *p* = 0.7710, unpaired *t*-test; Fig. 2D).

**Figure 2.**
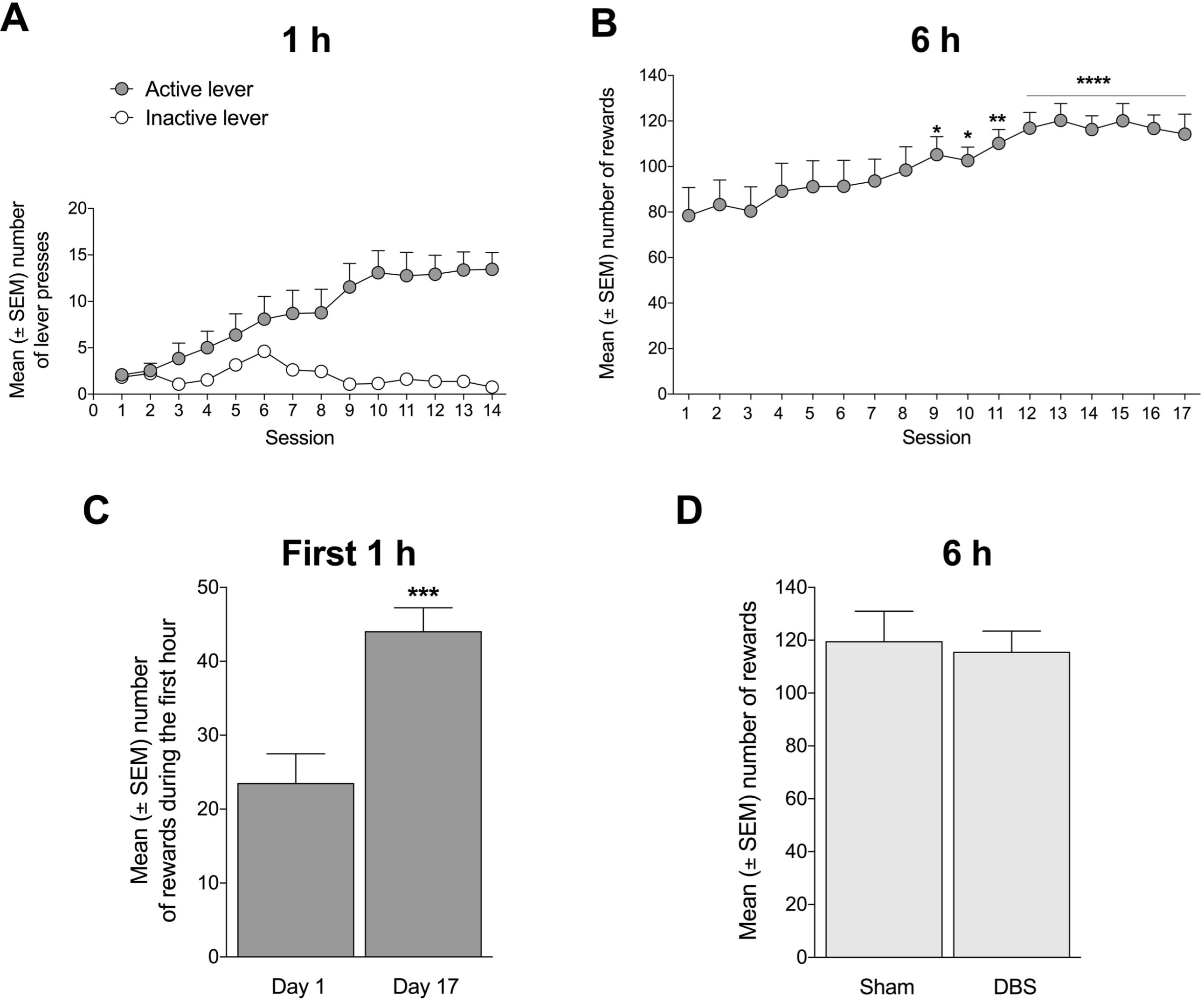
Training and escalation of cocaine self-administration. (A) Rats were first trained to self-administer cocaine (0.5 mg/kg/infusion) during 1-h sessions (*n* = 13). (B) The escalation of cocaine self-administration after seventeen 6-h sessions. \**p* < 0.05, \*\**p* < 0.01, \*\*\*\**p* < 0.0001, significantly different from day 1. (C) The first hour (i.e., the loading phase) of the 6-h self-administration session demonstrated significant escalation of cocaine intake during 17 sessions compared with day 1. \*\*\**p* < 0.001. (D) The rats were divided into sham and DBS groups according to equal baseline intake during the 6 h-session for the subsequent studies. *n* = 6-7. Note that the rats were not stimulated for the group assignment.

### Deep brain stimulation of the NAcc induces a modest increase in the escalation of cocaine self-administration

The rats were habituated by being connected to the stimulating cables for at least five sessions before being subjected to DBS treatment. After a stable level of self-administration was achieved (±10% over the last three sessions), the effect of DBS was studied for 4 consecutive days. The two-way repeated-measures ANOVA revealed a significant effect of treatment (*F*_1,6_ = 9.722, *p* = 0.0206; Fig. 3A) but no effect of time (*F*_1,6_ = 3.038, *p* = 0.1320) and no treatment × time interaction (*F*_1,6_ = 0.04698, *p* = 0.8356). When both sessions were averaged, DBS significantly increased cocaine self-administration (*t*_6_ = 3.118, *p* = 0.0206, paired *t*-test; Fig. 3B). Additionally, in the DBS group, a significant effect of treatment (*F*_1,6_ = 41.11, *p* = 0.0007; Fig. 3C) was found during the entire 6-h self-administration session but no effect of time (*F*_1,6_ = 1.726, *p* = 0.2369) and no treatment × time interaction (*F*_1,6_ = 1.903, *p* = 0.2170). When both sessions where averaged for the entire 6-h session, we found that DBS significantly increased cocaine self-administration (*t*_6_ = 6.411, *p* = 0.0007, paired *t*-test; Fig. 3D). No significant effect of treatment (*F*_1,5_ = 0.8128, *p* = 0.4086) or time (*F*_1,5_ = 0.004311, *p* = 0.9502) was observed in the Sham group during the first 90 min (Fig. 3E). Neither was there an effect of Sham-treatment when both sessions were averaged (*t*_5_ = 0.9015, *p* = 0.4086; Fig. 3F). No effect of treatment (*F*_1,5_ = 0.1045, *p* = 0.7596) or time (*F*_1,5_ = 0.005524, *p* = 0.9436) was observed during the entire 6-h self-administration session (Fig. 3G) or the average number of rewards during the 6-h session in the Sham-group (t_5_ = 0.3232, *p* = 0.7596; Fig. 3H).

**Figure 3.**
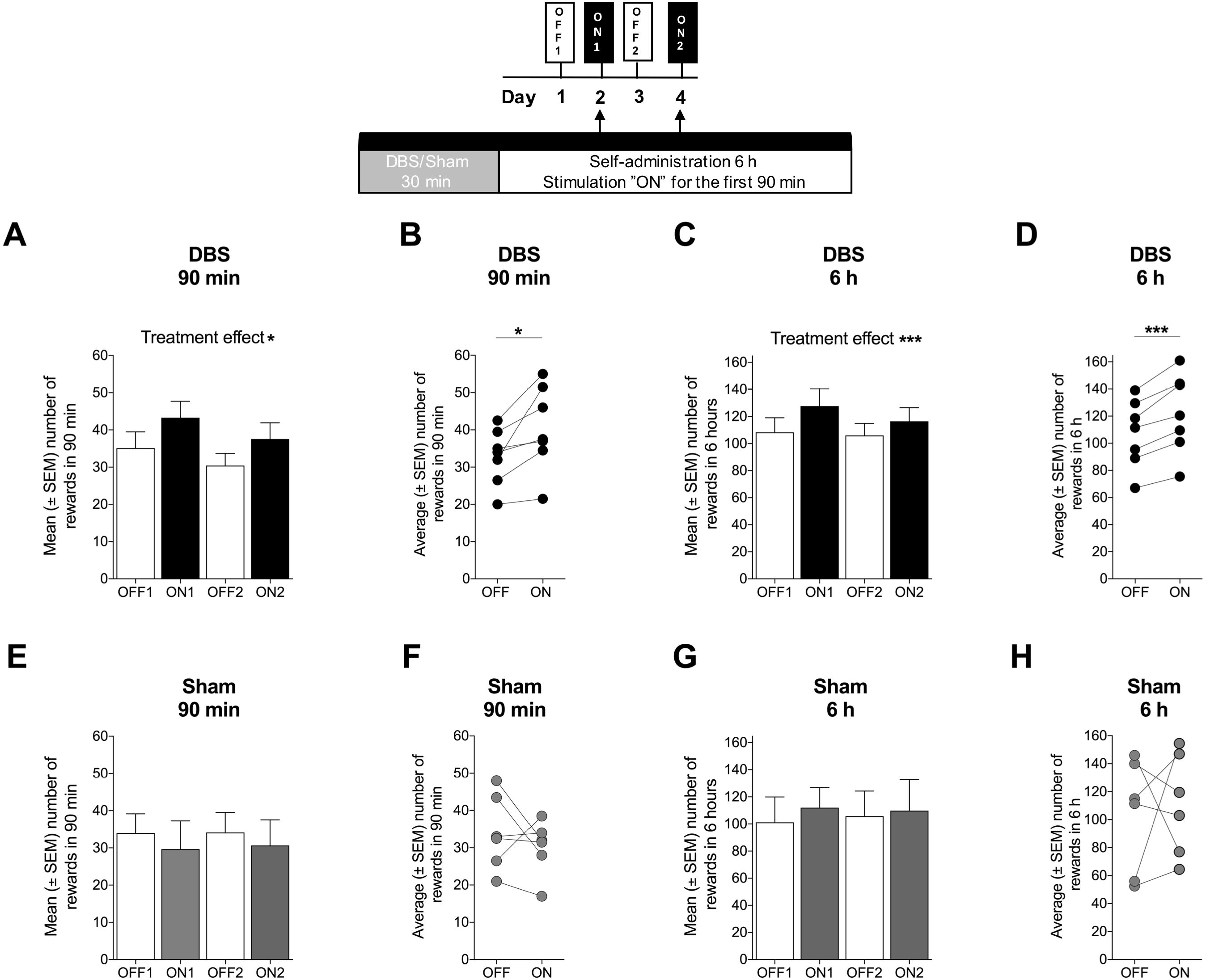
Deep brain stimulation of the NAcc shell increases cocaine self-administration. (A) Deep brain stimulation started 30 min before the self-administration session and remained on for a total of 2 h. An increase in cocaine intake during the first 90 min of the session was observed only in the first session of DBS (ON1). \**p* < 0.05. (B) Deep brain stimulation increased average cocaine intake (i.e., the average of both ON1 and ON2) during the first 90 min when DBS was on. \**p* < 0.05. (C) Deep brain stimulation increased cocaine intake during the entire 6-h session only in the first session of DBS. \**p* < 0.05. (D) Deep brain stimulation increased average cocaine intake (i.e., the average of ON1 and ON2) during the entire 6-h session. \*\*\**p* < 0.001. (E) Sham treatment had no effect on cocaine intake during the first 90 min of the 6-h session. (F) Sham treatment had no effect on average cocaine intake (i.e., the average of both ON1 and ON2) during the first 90 min of the self-administration session. (G) Sham treatment had no effect on cocaine intake during the entire 6-h session. (H) Sham treatment had no effect on average cocaine intake (i.e., the average of ON1 and ON2) during the entire 6-h session. *n* = 6-7.

### Deep brain stimulation of the NAcc increases cocaine-induced locomotor activity

Rats that received DBS immediately before the locomotor activity test exhibited an increase in behavioral sensitization to a intraperitoneal cocaine injection compared with the Sham group, indicated by a significant interaction in the two-way repeated-measures ANOVA (*F*_6,66_ = 2.645, *p* = 0.0232; Fig. 4A). Bonferroni’s multiple-comparison *post hoc* test did not reveal any significantly different time points between groups.

**Figure 4.**
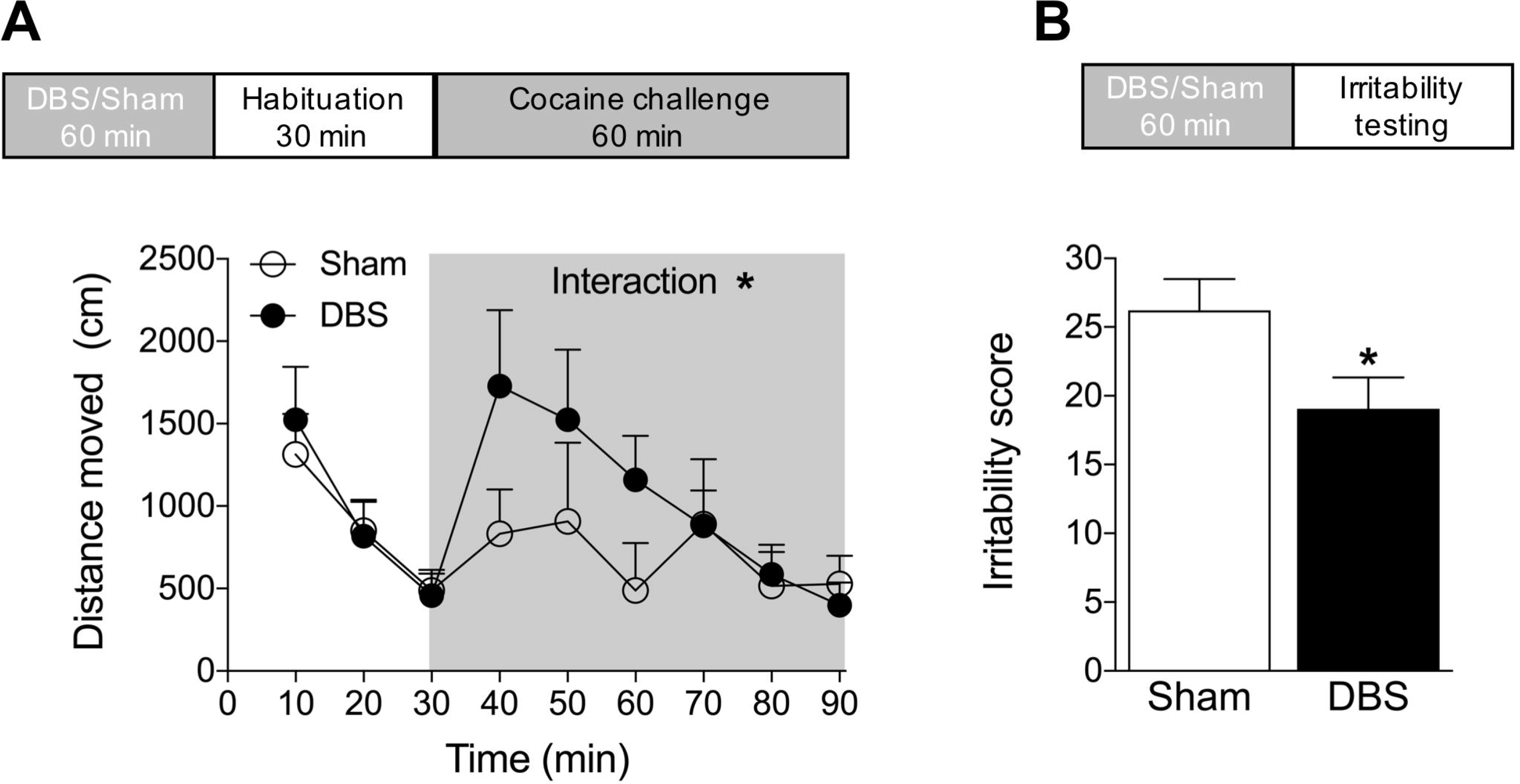
Deep brain stimulation of the NAcc shell increases the psychostimulant effect of cocaine but decreases irritability-like behavior. (A) Rats were exposed to DBS or sham treatment immediately before placing them in the open field. After 30 min of habituation, the rats were challenged with an acute injection of cocaine (10 mg/kg, i.p.). The DBS group exhibited an increase in cocaine-induced locomotor activity. \**p* < 0.05. *n* = 6-7. (B) Irritability-like behavior during cocaine withdrawal was decreased by DBS immediately before testing. \**p* < 0.05. *n* = 6-7.

### Deep brain stimulation of the NAcc decreases irritability-like behavior during cocaine withdrawal

Irritability-like behavior, a critical symptom of cocaine withdrawal in humans [42, 43], was measured 18 h into withdrawal after the escalation of cocaine self-administration. The correlation coefficient between blinded experimenters was high, ranging from *R* = 0.8845 (*p* < 0.0001) to *R* = 0.8963 (*p* < 0.0001). Deep brain stimulation that was applied immediately before behavioral testing decreased irritability scores (*t*_11_ = 2.204, *p* = 0.0497, Student’s two-tailed unpaired *t*-test; Fig. 4B).

### Deep brain stimulation of the NAcc shell increases GluR1 levels in the CeA

Following 60 min of DBS or sham treatment, we measured the levels of the AMPA receptor subunits GluR1/GluA1 and GluR2/GluA2 in both synaptosomal and cytosolic fractions of eight different brain regions: NAcc shell and core, dorsal striatum, vPFC, dPFC, CeA, insula, and VTA. In the synaptosomal fractions, an increase in GluR1 levels was found only in the CeA (*p* = 0.033, Bonferroni’s *post hoc* test; Fig. 5A). No significant changes in GluR2 were found in the synaptosomal fractions (*p* > 0.4 for all analyzed regions, Bonferroni’s *post hoc* test; Fig. 5B). In the cytosolic fractions, a significant increase in GluR1 levels was found only in the CeA (*p* = 0.007, Bonferroni’s *post hoc* test; Fig. 5C). No significant changes in GluR2 were found in the cytosolic fractions, although a trend toward an increase in the CeA was observed in the DBS group (*p* = 0.094, Bonferroni’s *post hoc* test; Fig. 5D).

**Figure 5.**
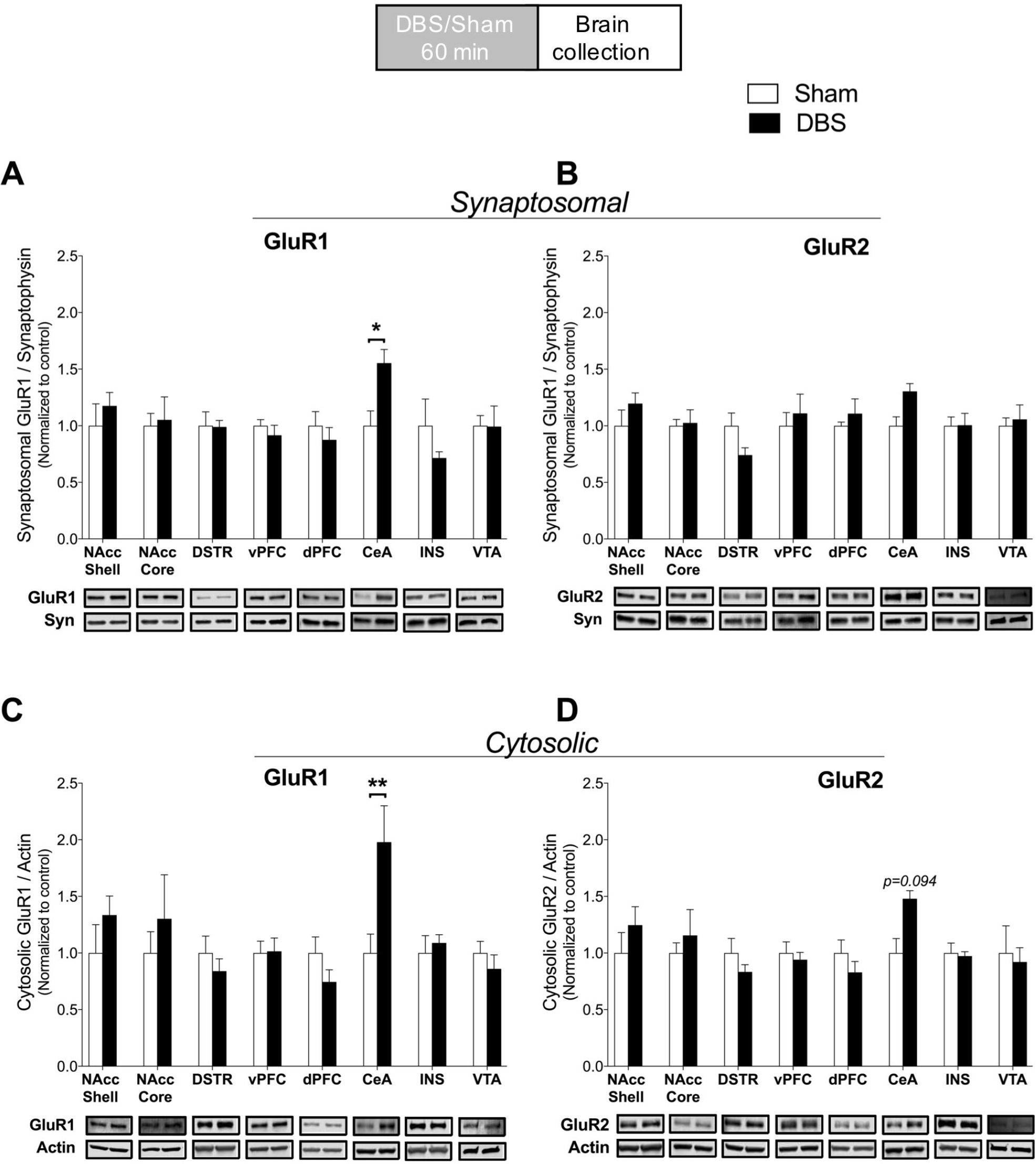
Deep brain stimulation increases GluR1/GluA1 levels in the central nucleus of the amygdala (CeA) in both synaptosomal and cytosolic fractions. (A) In the synaptosomal fractions of eight brain regions, DBS increased the levels of GluR1 in the CeA (*p* = 0.033, Bonferroni *post hoc* test). (B) No changes in GluR2/GluA2 were observed in any of the studied brain regions. (C) In the cytosolic fractions, DBS increased the levels of GluR1 in the CeA (*p* = 0.007, Bonferroni *post hoc* test). (D) No significant changes in GluR2 were observed, although a trend was found in the CeA (*p* = 0.094, Bonferroni *post hoc* test). NAcc, nucleus accumbens; DSTR, dorsal striatum; vPFC, ventral prefrontal cortex; dPFC, dorsal prefrontal cortex; CeA, central nucleus of the amygdala; INS, insular cortex; VTA, ventral tegmental area. The data are expressed as mean ± SEM. Representative Western blots are shown below the figures. The approximate molecular weights of the observed band sizes were ~100 kDa for GluR1 and GluR2, ~38 kDa for synaptophysin (Syn), and ~42 kDa for β-actin (Actin). *n* = 5-7/group. \**p* < 0.05, \*\**p* < 0.01.

## Discussion

Contrary to our hypothesis, the present study demonstrated that high-frequency DBS of the NAcc shell did not decrease the escalated cocaine intake but instead actually produced a slight increase in intake and in cocaine-induced locomotor activity in cocaine-dependent rats, without affecting baseline exploratory activity during habituation. Interestingly, DBS decreased rather than increased irritability-like behavior. Finally, DBS of the NAcc shell increased both synaptosomal and cytosolic levels of GluR1 specifically in the CeA, with no effect on GluR2.

Previous studies found that DBS of the NAcc shell but not the core attenuated the reinstatement of cocaine intake in nondependent rats [21, 22]. However, DBS was later found to also attenuate the reinstatement of sucrose seeking in rats, suggesting that the effect of DBS generalizes to both drug and natural reinforcers, thus questioning its applicability in the treatment of addiction [44]. Nonetheless, to our knowledge, only one previous study has investigated the effect of DBS on cocaine self-administration itself, in which no effect of either low-frequency (20 Hz) or high-frequency (160 Hz) stimulation on cocaine intake was observed in nondependent rats during the 90 min self-administration session [45]. In addition, a recent study found that optogenetic stimulation of the NAcc shell had no effect on the motivation to self-administer cocaine and reduced cocaine-seeking behavior during extinction [46], which is consistent with the findings of Vassoler and colleagues. Interestingly, however, the same study also found that selective optogenetic stimulation of the NAcc shell-to-lateral hypothalamus pathway enhanced the motivation to self-administer cocaine and increased context-induced cocaine-seeking behavior. Therefore, the increase in cocaine self-administration that was observed in the present study may be mediated by the stimulation of this γ-aminobutyric acid (GABA)ergic projection from the NAcc shell to the lateral hypothalamus that mediates motivated behaviors. Consistent with the increase in cocaine self-administration, the present study also found that DBS of the NAcc shell sensitized the rats to the psychostimulant effect of cocaine. In contrast to our results, high-frequency DBS of the NAcc shell transiently suppressed locomotor sensitization to cocaine in mice [47]. This discrepancy may be explained by the fact that we used cocaine-dependent rats, whereas Creed et al (2015) studied cocaine sensitization in mice that received only seven injections of cocaine. We also found that DBS of the NAcc shell immediately before testing decreased rather than increased irritability-like behavior during cocaine withdrawal. Together with anxiety and dysphoria, irritability-like behavior is a key negative emotional state that characterizes the withdrawal syndrome in humans [48]. Irritability-like behavior is also a critical symptom of cocaine withdrawal in humans [42, 43]. Such a decrease in irritability-like behavior may contribute to the lower motivation to seek cocaine that is observed in abstinent rats after DBS [22]. Thus, DBS of the NAcc may have two opposite effects, depending on whether cocaine is present or not. Deep brain stimulation of the NAcc shell may decrease craving [44] and negative affective states (present study) during abstinence when cocaine is unavailable and may increase the psychostimulant effect of cocaine and amount of cocaine that is self-administered when cocaine is present (present study).

Glutamate signaling in the brain, especially in the PFC, NAcc, VTA, and amygdala, plays an important role in molecular and behavioral plasticity that is associated with addictive drugs [49]. Drugs of abuse, including morphine, cocaine, and alcohol, have been found to elevate levels of the ionotropic AMPA-selective GluR1 subunit in the VTA and NAcc [50–54]. Morphine self-administration has been found to increase GluR1 receptor labeling in the basolateral amygdala (BLA) but not CeA [55]. Furthermore, the dynamic regulation of AMPA receptors during cocaine self-administration has been suggested to facilitate subsequent use [23]. Importantly, the incubation of cocaine craving was found to be caused by an increase in GluR1-containing/GluR2-lacking calcium-permeable AMPA receptors in the NAcc [24]. Additionally, AMPA receptor blockade has been found to attenuate cocaine-seeking behavior [56–58], whereas AMPA receptor agonism promotes cocaine reinstatement [25–27]. AMPA receptors exist both at synapses and in internal stores and undergo trafficking [39–41]. Thus, in the present study, we measured both synaptosomal and cytosolic levels of GluR1 and GluR2. Interestingly, we found that DBS of the NAcc shell induced a robust increase in both synaptosomal and cytosolic levels of GluR1 in the CeA. Although the levels of cytosolic GluR2 in the CeA showed a similar trend as GluR1, these changes were not significant after Bonferroni correction. The fact that DBS increased GluR1 levels in both the synaptosomal and cytosolic fractions suggests regulation at the translational or even transcriptional level rather than at the trafficking level between cytosolic and synaptic compartments. Interestingly, DBS of the PFC has also been found to increase GluR1 levels, although these changes were seen in the NAcc core and shell [59], which may be expected given the glutamatergic projections from the PFC to the NAcc. The present findings with regard to glutamate receptors suggest that DBS stimulation of the NAcc can affect GluR1 levels in the amygdala, possibly via antidromic stimulation. The BLA has glutamatergic afferents to the NAcc, mainly to the core, that are considered key pathways for the expression of motivated behavior that is driven by emotionally and motivationally relevant stimuli [60, 61]. Nonetheless, the exact role of higher GluR1 levels in the CeA in the promotion of drug self-administration remains to be investigated.

It is important to note that in this study rats underwent several stimulating sessions in the course of the experiment. Therefore, it is possible that there is accumulating effects and/or gradual adaptation to the effects of DBS. Importantly, however, DBD treatment in humans is also given repeatedly and thus, it is also relevant to study the effects of repeated DBS in a preclinical model of cocaine addition.

Both preclinical and clinical studies that have validated the safety and efficacy of DBS as a treatment for substance use disorder have to date focused on the NAcc as a therapeutic target. This therapeutic strategy needs to be optimized, with a better understanding of the specific mechanisms by which DBS of the NAcc modulates drug craving and possibly intake [15, 16]. Although the attenuation of cocaine-induced reinstatement by DBS of the NAcc shell has been previously reported, the present study does not support the NAcc shell as a therapeutic target of high-frequency DBS in the treatment of cocaine addiction. Patients with severe cocaine use disorder have a high probability of lapses and relapses, and such lapses may be exacerbated by DBS of the NAcc shell. Thus, investigating other parameters of stimulation (e.g., low-frequency DBS that is refined by dopamine D_1_ receptor inhibition) would be worthwhile [47, 62]. Addittionally, the effects of DBS of the NAcc core should be further explored as well as other target regions of DBS, such as the PFC, habenula, and subthalamic nucleus (STN)[63], for the treatment of addiction [64, 65] may yield better results. Indeed, converging evidence suggests that the STN may be a key target for DBS in the treatment of addiction [32, 66, 67]. Additionally, stimulation of the lateral habenula has been shown to decrease cocaine-seeking behavior [68] and alcohol consumption [69].

## Conclusions

In the present study we found that high-frequency DBS of the NAcc shell did not decrease the escalation of cocaine intake but instead slightly increased cocaine intake and cocaine-induced locomotion in cocaine-dependent rats accompanied with a robust increase in both synaptosomal and cytosolic levels of GluR1 in the CeA. These preclinical findings do not support the use of high-frequency DBS of the NAcc shell as a therapeutic approach to decrease intake in cocaine dependent subjects and therefore questions its suitability for the treatment of cocaine addiction. However, we confirmed a beneficial effect of DBS of the NAcc on irritability-like behavior during abstinence, suggesting that DBS may still ease withdrawal and possibly craving during abstinence.

## Acknowledgements

We thank Gustav Montenegro for proofreading the manuscript and Dr. Fair Vassoler for technical advice.

